# Accurate *de novo* identification of biosynthetic gene clusters with GECCO

**DOI:** 10.1101/2021.05.03.442509

**Authors:** Laura M. Carroll, Martin Larralde, Jonas Simon Fleck, Ruby Ponnudurai, Alessio Milanese, Elisa Cappio, Georg Zeller

**Affiliations:** Structural and Computational Biology Unit, EMBL, Heidelberg, Germany; Department of Biology, ETH Zürich, Zürich, Switzerland; Present address: Department of Biosystems Science and Engineering, ETH Zürich, Basel, Switzerland

## Abstract

Biosynthetic gene clusters (BGCs) are enticing targets for (meta)genomic mining efforts, as they may encode novel, specialized metabolites with potential uses in medicine and biotechnology. Here, we describe GECCO (GEne Cluster prediction with COnditional random fields; https://gecco.embl.de), a high-precision, scalable method for identifying novel BGCs in (meta)genomic data using conditional random fields (CRFs). Based on an extensive evaluation of *de novo* BGC prediction, we found GECCO to be more accurate and over 3x faster than a state-of-the-art deep learning approach. When applied to over 12,000 genomes, GECCO identified nearly twice as many BGCs compared to a rule-based approach, while achieving higher accuracy than other machine learning approaches. Introspection of the GECCO CRF revealed that its predictions rely on protein domains with both known and novel associations to secondary metabolism. The method developed here represents a scalable, interpretable machine learning approach, which can identify BGCs *de novo* with high precision.

## INTRODUCTION

Host-associated and environmental microbes alike are capable of producing a wide array of secondary metabolites through which they interact with their environments.^1^ These metabolites equip their producer with a chemical repertoire to respond to stressors, which may confer competitive advantages over other organisms in their environmental niche.^2,3^ In human host-associated microbial communities, secondary metabolites can also modulate host health via a range of processes, including immune system regulation, xenobiotic and nutrient metabolism, and cancer susceptibility/resistance.^3,4^ Beyond their natural purposes, many microbial secondary metabolites have found important uses in medicine, including as first-in-class antimicrobial, anticancer, and antidiabetic drugs.^1,5,6^

Due to the biomedical and biotechnological interest in microbial secondary metabolites, there is a strong incentive to identify novel natural products. Genome mining efforts have successfully made use of the fact that a large proportion of the enzymatic pathways responsible for secondary metabolite production are encoded by physically clustered groups of genes called biosynthetic gene clusters (BGCs).^1,7–9^ Recently, the development of computational tools for BGC detection has been further fueled by the ever-increasing availability of microbial genomic and metagenomic data.^7–8^ Currently, *in silico* methods used to identify BGCs in (meta)genomic sequencing data can largely be categorized into two groups. “Rule-based” approaches (e.g., antiSMASH, PRISM)^8,11,12^ use hard-coded BGC detection “rules” to identify BGCs in (meta)genomic data based on signature genes.^9^ These approaches display a high degree of precision (i.e., low false positive rates) but are unable to detect novel BGCs of unknown architecture. To prioritize the detection of novel BGCs, “model-based” approaches have been developed.^7,9^ The most widely used representative of this group, ClusterFinder, relies on a hidden Markov model (HMM) to segment (meta)genomic sequences into BGC and non-BGC regions based on local enrichment of protein domains characteristic of biosynthetic genes.^7^ More recently, DeepBGC, which employs a three-layer Bidirectional Long Short-Term Memory (BiLSTM) recurrent neural network (RNN)^13^, was shown to yield more accurate *de novo* BGC predictions than HMM-based ClusterFinder.^9^

Conditional random fields (CRFs) are an alternative machine learning (ML) approach to HMMs and BiLSTMs for sequence segmentation. These discriminative graphical models (Fig. 1 and Supplementary Figure S1) have been shown to outperform generative models, such as HMMs, in various application domains.^14,15^ Furthermore, compared to their “black box” RNN counterparts, CRFs have the advantage of being inherently interpretable, an important feature in a biomedical context.^16^ Here, we describe GECCO (GEne Cluster prediction with COnditional random fields; https://gecco.embl.de), a high-precision, scalable method for *de novo* BGC identification in microbial genomic and metagenomic data. On the basis of a newly developed, extensive *de novo* BGC prediction benchmarking framework, we show that GECCO is not only more accurate than state-of-the-art *de novo* BGC detection approaches, but also more computationally efficient. As an interpretable ML model, GECCO can moreover provide insights into BGC biology, architecture, and function.

**Figure 1.**
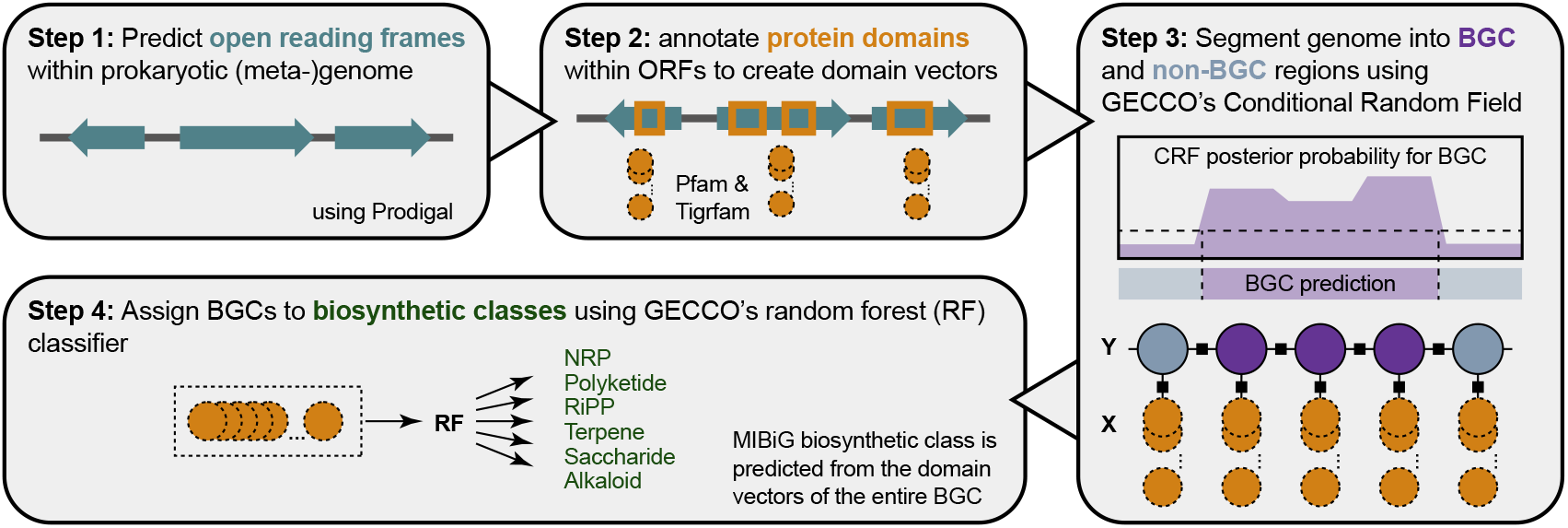
Graphical depiction of the biosynthetic gene cluster (BGC) identification and classification approach developed here and implemented in GECCO (GEne Cluster prediction with COnditional random fields). Briefly, GECCO identifies open reading frames (ORFs) in an assembled prokaryotic (meta)genome (Step 1). Protein domains are annotated in the resulting ORFs using profile hidden Markov models (pHMMs; Step 2). The resulting ordered domain vectors are treated as features, and a conditional random field (CRF) is used to predict whether each feature belongs to a BGC or not (Step 3). Predicted BGCs are classified into one of six major biosynthetic classes as defined in the Minimum Information about a Biosynthetic Gene cluster (MIBiG) database using a Random Forest classifier (Step 4).

## RESULTS

### GECCO: a CRF-based *de novo* BGC detection tool

To train a CRF that could identify novel BGCs in (meta)genomic sequences, a training/cross-validation (CV) data set was constructed by embedding known BGCs into long, BGC-negative fragments of prokaryotic genomes (Fig. 1 and Supplementary Figures S1 and S2). Briefly, known BGCs present in the Minimum Information about a Biosynthetic Gene cluster (MIBiG) database^1^ were embedded into randomly selected prokaryotic contigs, in which other known and predicted BGCs had been masked (see section “Data acquisition and feature construction” below). To construct the feature matrix for training, open reading frames (ORFs) were identified and annotated with protein domains, using one of fourteen combinations of databases in which protein families are represented by profile hidden Markov models (pHMMs; see Fig. 1, section “Data acquisition and feature construction” below, and Supplementary Figures S1 and S2). As the protein family resources are broader in scope than what may be needed for BGC identification and their combinations potentially redundant, an additional feature selection approach was implemented in GECCO: to identify domains that are either most strongly enriched or depleted in BGCs, we nested a two-sided Fisher’s Exact Test (FET) into the CV employed for fitting GECCO’s CRF. Within each CV fold, we iteratively retrained GECCO using only the top domains associated with BGC presence or absence to estimate how far the CRF feature space (i.e. the domain pHMMs used for annotation) could be reduced to gain speed while retaining optimal prediction accuracy (Supplementary Figures S3 and S4a).

### GECCO provides superior precision and speed relative to state-of-the-art *de novo* BGC prediction methods

To construct a benchmark data set in a way that guarantees that training and test data are disjoint, we partitioned MIBiG v2.0^17^ into (i) BGCs for training that were already contained in an earlier MIBiG version (v1.3, which was also originally used to train DeepBGC and DeepBGC’s re-trained implementation of ClusterFinder)^9^, and (ii) selected BGCs for testing that were newly added in subsequent updates of MIBiG; from this test set we also removed BGCs that were very similar in architecture to any instance contained in MIBiG v1.3 (see section “Data acquisition and feature construction” below for additional details). This yielded a final test set of 376 prokaryotic contigs which each had an embedded BGC that was exclusively present in MIBiG v2.0 (referred to hereafter as the “376-genome test set”). We also used two additional, previously constructed test sets containing thoroughly annotated genomes of well-studied BGC producer organisms (the “six genome test set” used by Hannigan, et al.^9^ to evaluate DeepBGC and the “nine genome bootstrapping set” used by Hannigan, et al.^9^ for hyperparameter tuning, validation, and testing of DeepBGC) and removed all instances of BGCs similar to those in these additional test sets from the MIBiG v1.3-based training set. To ensure a fair comparison of BGC detection methods, we retrained DeepBGC and GECCO on this very same training set, using BGCs from MIBiG v1.3 which were absent from all test sets. We evaluated the performance of both methods using (i) the 376-genome test set (the main test set presented in this study; Fig. 2a-c), as well as (ii) the six- and (iii) nine-genome test sets^9^ and (iv) 10-fold CV using BGCs from MIBiG v1.3 (Supplementary Figures S4-S6). We additionally compared GECCO and the re-trained implementation of DeepBGC to the original DeepBGC, as well as DeepBGC’s “original” and “re-trained” implementations of the ClusterFinder algorithm (also trained on BGCs from MIBiG v1.3), using the three aforementioned test sets (Fig. 2a-c and Supplementary Figure S6).^7,9^

**Figure 2.**
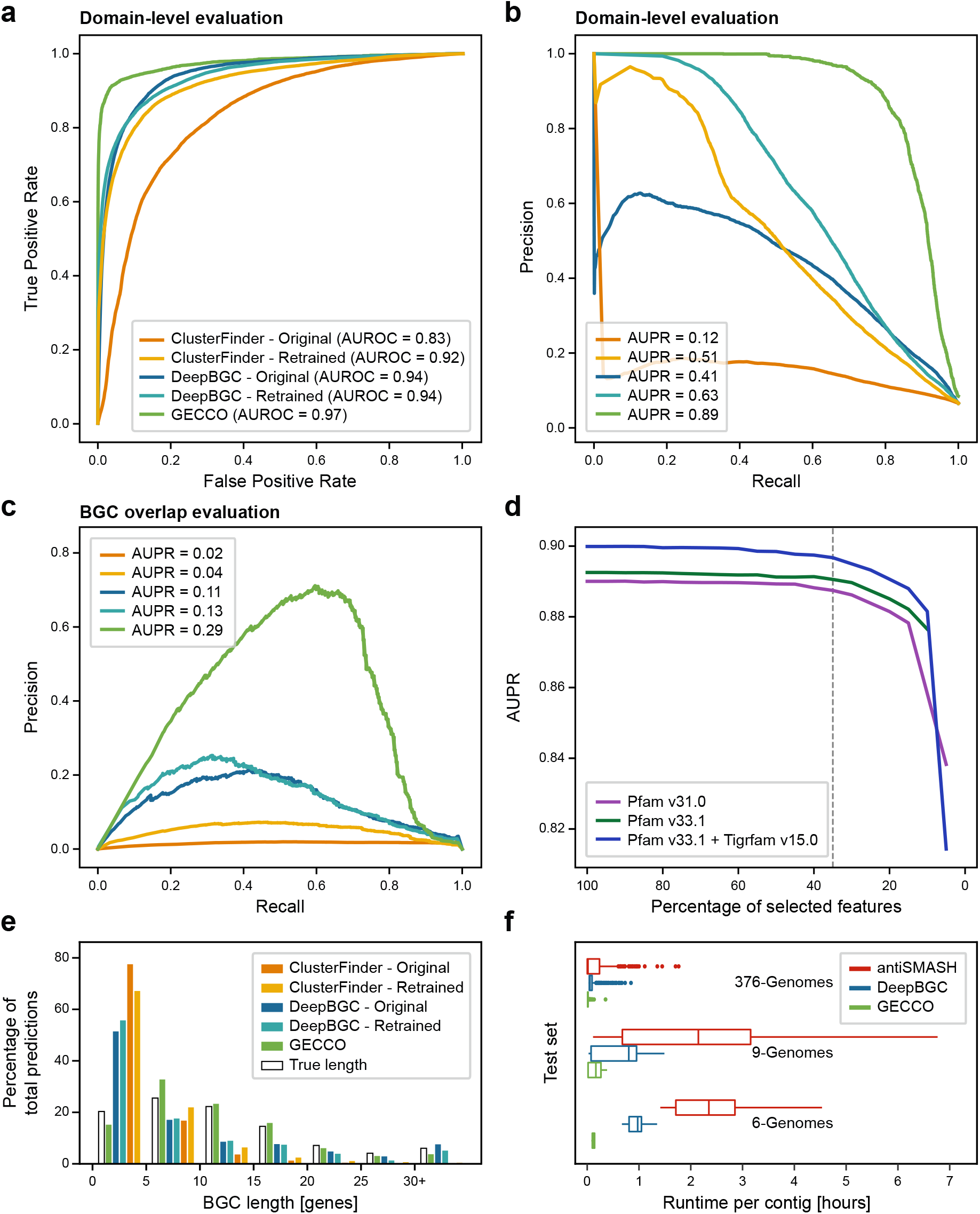
**(a)** Domain-level receiver-operating characteristic (ROC) curves, **(b)** domain-level precision-recall (PR) curves comparing original and retrained implementations of ClusterFinder (ClusterFinder-Original and ClusterFinder-Retrained, respectively) and DeepBGC (DeepBGC-Original and DeepBGC-Retrained, respectively) with GECCO (trained on a subset of Pfam v33.1 and Tigrfam v15.0 domains). **(c)** PR curves calculated from segment overlap (>50%) of predicted and known BGCs (see Supplementary Figures S4 and S7). All models (a-c) were trained on BGCs from MIBiG v1.3, evaluated on BGCs from MIBiG v2.0 not contained in v1.3 (i.e., the 376-genome test set); area under the curve (AUROC and AUPR) values are reported in legends. **(d)** AUPR values (Y-axis) versus percentage of Fisher’s Exact Test-selected features (*T*; X-axis) included in CRFs trained on BGCs from MIBiG v2.0, using domains from several (combinations of) databases (see inset). The default value of *T* chosen for GECCO is denoted by the dashed line (*T =* 0.35). **(e)** Histogram of predicted BGC lengths (in number of genes; X-axis) relative to true lengths among genomes in the 376-genome test set. The Y-axis denotes the percentage of total BGC predictions for each method. (f) Runtime per contig required to detect and classify BGCs in each test set using antiSMASH, DeepBGC, and GECCO.

For direct comparability to previous evaluations,^9^ we first conducted a receiver operating characteristic (ROC) analysis on the level of individual domains, i.e. based on a per-domain assessment of true positives, true negatives, false positives and false negatives (Fig. 2a, Supplementary Figure S4a). Based on area under (AU) the ROC curve values, GECCO showed superior performance compared to all DeepBGC/ClusterFinder implementations on the 376-and six-genome test sets (Fig. 2a and Supplementary Figure S6) and during 10-fold CV (GECCO AUROC = 0.97; Supplementary Figure S5). On the nine-genome test set, GECCO and the original implementation of DeepBGC performed equally (AUROC = 0.94; Supplementary Figure S6). From the same true/false positive/negative metrics, we also constructed per-domain precision-recall (PR) curves (Fig. 2b, Supplementary Figure S4a). These evaluations showed superior performance of GECCO compared to all DeepBGC/ClusterFinder implementations for all three test sets (Fig. 2b and Supplementary Figure S6) and during 10-fold CV (GECCO AUPR = 0.73; Supplementary Figure S5). In addition to evaluating model performance at the domain level, we also assessed to which extent predicted BGCs overlapped with known BGCs by calculating precision and recall from true and false positive BGC segments, as well as false negative non-BGC segments, for each model (referred to hereafter as the “segment overlap” metric; Supplementary Figure S4b). Based on PR curves constructed from this segment overlap metric, GECCO achieved substantially higher AUPR than all implementations of DeepBGC/ClusterFinder (Fig. 2c and Supplementary Figures S4-S7). These evaluations demonstrate that GECCO is capable of detecting BGCs *de novo* with unprecedented accuracy, primarily by more precisely locating their boundaries; this also greatly alleviates the problem of fragmented predictions, which other methods suffer from (Fig. 2e).

We moreover used the training data to optimize GECCO’s feature space. We found that feature inclusion thresholds *T*= [35,100] (percentage of retained domain features) achieved highly similar AUPR and *F*_1_ scores (Fig. 2d and Supplementary Figure S3), suggesting that 65% of features can be discarded without noticeable sacrifices in accuracy. Among the domain resources used for feature generation, a combination of TIGRFAM v15.0^18^ and Pfam v33.1^19^ with *T =* 35% achieved among the highest AUROC and AUPR scores, and was thus chosen as the final model for BGC detection in GECCO (10-fold CV AUROC = 0.96, AUPR = 0.89; Fig 2d, Supplementary Figure S3). To explore GECCO’s ability to identify novel BGC classes not currently represented in MIBiG, leave-one-type-out (LOTO) CV was used. In LOTO, one biosynthetic class of BGCs is completely removed from the training set during CV to specifically assess its re-discovery in the test set. GECCO achieved LOTO AUROC scores > 0.98 for four of six classes and 0.91 and 0.88 AUROC for MIBiG’s ribosomally synthesized and post-translationally modified peptide (RiPP) and Saccharide classes, respectively (Supplementary Figure S8). To determine the MIBiG biosynthetic class for each newly predicted BGC, a separate random forest (RF) classifier was trained and evaluated using five-fold CV, as has been previously proposed^9^ (see section “Prediction of biosynthetic class” below; Fig. 1). Using the domain composition associated with each BGC as features, the RF classifier achieved AUROC scores > 0.90 for all classes (Supplementary Figure S9).

In a final benchmark, we compared the runtime between GECCO, DeepBGC, and antiSMASH using the three test sets, as well as all representative genomes in the proGenomes2 database, a comprehensive resource of prokaryotic genome sequences (containing 627,182 contigs from 12,221 genomes; Fig. 2f and Supplementary Figure S10).^20^ Using a single CPU, GECCO was over three times faster than both antiSMASH and DeepBGC (Fig. 2f and Supplementary Figure S10).

Taken together, this comparative evaluation, to our knowledge, is the most comprehensive benchmarking of *de novo* BGC prediction tools conducted to date. It clearly demonstrates that GECCO greatly improves the accuracy of *in silico* BGC identification over the state of the art, while also being computationally efficient.

### GECCO’s CRF-based approach provides insight into the biosynthetic potential of microbes

To compare GECCO BGC predictions to those produced by other tools on a real-world data set, each of GECCO, DeepBGC, and antiSMASH were used to identify and classify BGCs among all 12,221 representative genomes in the proGenomes2 database. Notably, the majority of BGCs predicted by either GECCO or DeepBGC were not detected using the rule-based approach implemented in antiSMASH (*n =* 59,041 antiSMASH BGCs using default parameters; Fig. 3ab). Overall, GECCO predicted nearly twice as many BGCs as antiSMASH, but far fewer than DeepBGC (*n =* 115,131 and 470,137 GECCO and DeepBGC BGCs, respectively), consistent with the above evaluations showing a clear tendency of GECCO to produce fewer false positives and fragmented predictions (Fig. 2a-c,e and Supplementary Figures S4-S7).

**Figure 3.**
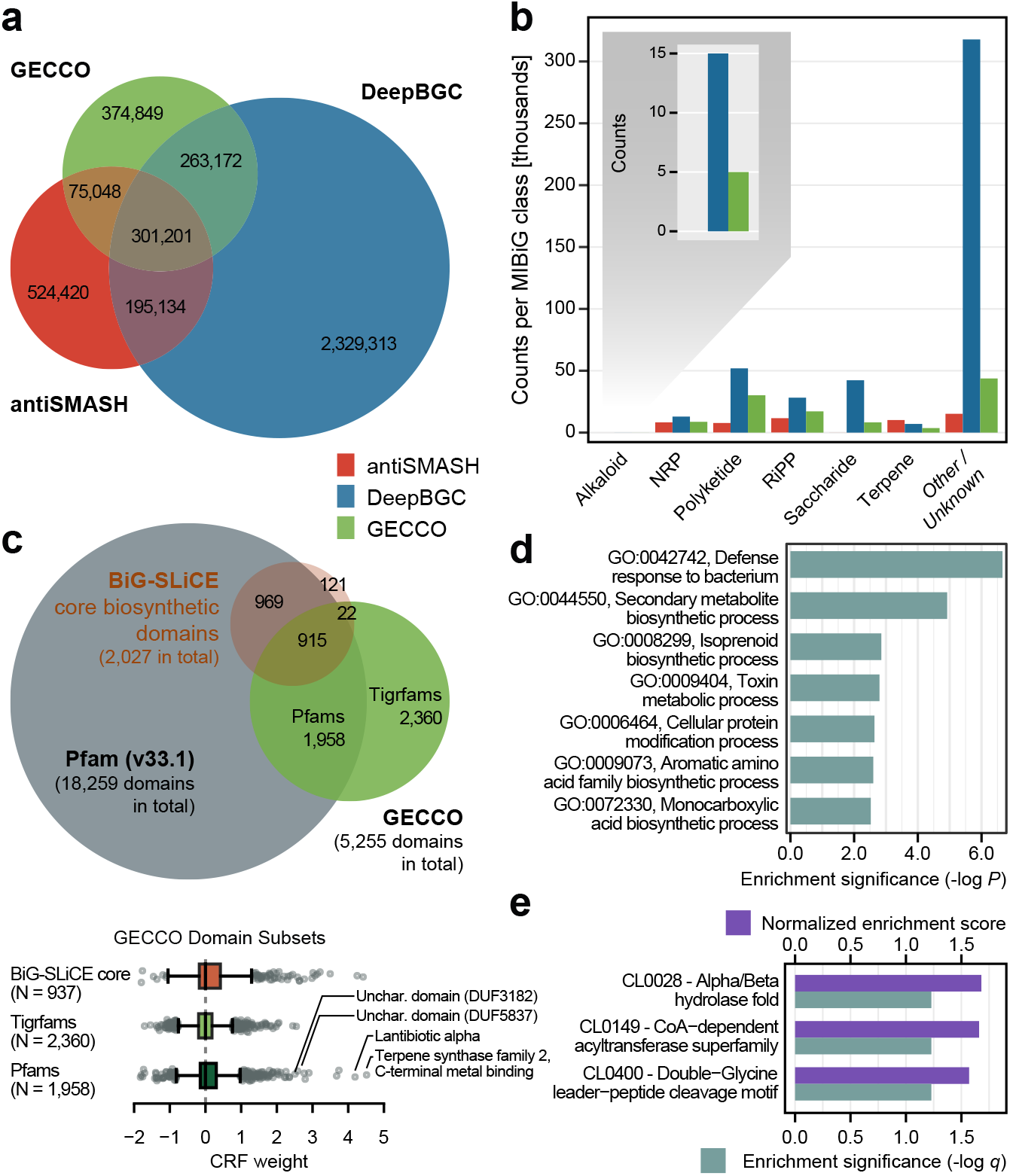
**(a)** Venn diagram of biosynthetic gene cluster (BGC) overlap, constructed using the presence and absence of individual genes in BGCs identified in 12,221 representative microbial genomes available in the proGenomes2 database using each of antiSMASH, DeepBGC, and GECCO. If a gene was contained within BGC predictions of more than one method, it was counted in the respective intersection area. **(b)** Predicted MIBiG biosynthetic classes (X-axis) associated with BGCs identified in the same 12,221 genomes using each of antiSMASH, DeepBGC, and GECCO. The Y-axis denotes the number of BGCs assigned to a given biosynthetic class. BGCs assigned to multiple classes are omitted. **(c)** Venn diagram and boxplots of GECCO CRF weights (X-axis), constructed using protein domains used by (i) GECCO, (ii) BiG-SLiCE, and (iii) and Pfam v33.1. GECCO domains were derived from either Pfam v33.1 or Tigrfam v15.0 and were selected based on their association with BGC presence/absence using Fisher’s Exact Test (FET) and an FET-inclusion threshold (*T*) of 35% (*T =* 0.35). BiG-SLiCE domains correspond to those present in the core biosynthetic domain set used by BiG-SLiCE v1.1.0. (d) Top Gene Ontology (GO) terms (Y-axis) enriched in BGCs, obtained using the Kolmogorov-Smirnov test/weight01 algorithm implemented in topGO (enrichment significance > 2.75). (e) Pfam clans (Y-axis) enriched in BGCs (X-axis; false discovery rate [FDR]-adjusted *P <* 0.10). Normalized Enrichment Scores (NES) were obtained using the fgsea R package. For (d and e), enrichment significance values correspond to the negated base-10 logarithm of each term’s *P*-value.

To investigate which protein domains GECCO relied on for BGC detection, we first analyzed which protein domains were retained in the first feature elimination step. Notably, nearly half of all GECCO protein domains (2,382 of 5,255 total GECCO protein domains, 45.3%) were derived from the TIGRFAM database (Fig. 3c), highlighting the complementary nature of the Pfam and TIGRFAM databases for BGC prediction optimization. When compared to a collection of protein domains previously associated with secondary metabolism (used by BiG-SLiCE v1.1.0)^21^, nearly half of these core biosynthetic domains were included in GECCO’s model (937 of 2,027 BiG-SLiCE core domains, 46.2%; Fig. 3c). Domains in the core biosynthetic set/GECCO intersection received more positive (i.e., BGC-associated) CRF weights relative to TIGRFAM domains not present in the core biosynthetic domain set, but not relative to Pfam domains not present in the core biosynthetic domain set (two-sided Mann-Whitney *U* test raw *P =* 3.12e-07 and 0.10, respectively; Fig. 3c). However, many other domains outside of the comparatively small core biosynthetic space received CRF weights with comparably high (absolute) values. Domains with negative weights were important for capturing non-BGC regions (Fig. 3c); however, some of the most highly weighted (i.e., BGC-associated) domains were not members of the core biosynthetic set (Fig. 3c, Supplementary Table S1). Among these (CRF weight > 4.0) were (i) terpene synthase family 2, C-terminal metal binding domain PF19086 and (ii) lantibiotic alpha domain PF14867, both of which have previously been associated with secondary metabolite production (Fig. 3c, Supplementary Table S1). Interestingly, among the highest-weighted, BGC-associated domains (CRF weight > 2.0) that were not members of the core biosynthetic set were three domains of unknown function (DUF): (i) PF19155 (DUF5837), which is associated with a cyanobactin (RiPP) BGC, tenuecyclamide A (MIBiG ID BGC0000480); (ii) PF11379 (DUF3182), a Proteobacteria-restricted protein of unknown function (InterPro ID IPR021519); (iii) PF17537 (DUF5455), a protein of unknown function found in Proteobacteria, which contains three predicted trans-membrane regions (InterPro ID IPR035210; Supplementary Table S1). Their importance for BGC prediction with GECCO suggests that functional studies of these domains in the context of secondary metabolism are warranted.

To be able to observe coherent biological functions among the domain weights learned by the GECCO CRF, beyond the most strongly associated domains, we used Gene Ontology (GO)^22^ and Pfam (structural) clan^19^ annotations. This led to the identification of 33 biological processes (BPs) and 22 molecular functions (MFs) enriched in BGCs (topGO Kolmogorov-Smirnov *P <* 0.05), with “defense response to bacterium” (GO:0042742), “secondary metabolite biosynthetic process” (GO:0044550), “isoprenoid biosynthetic process” (GO:0008299), and “toxin metabolic process” (GO:0009404) showcasing the strongest associations (all BGC enrichment scores > 2.5; Fig. 3d and Supplementary Figure S11). Three Pfam clans were additionally enriched in BGCs (false discovery rate-corrected *P <* 0.10): Alpha/Beta hydrolase fold (CL0028), CoA-dependent acyltransferase superfamily (CL0149), and Double-Glycine leader-peptide cleavage motif (CL0400; Fig. 3e and Supplementary Figure S12). Collectively, these results indicate that the GECCO CRF relies on domains associated with secondary metabolite production for BGC inference.

## DISCUSSION

ML approaches have revolutionized numerous disciplines and are being increasingly employed to solve problems in biological and medical realms.^23–25^ Models that can account for sequential data are particularly attractive when leveraging genomic data to make predictions, as feature context and order (e.g., for genes, domains) may be important.^9^ CRFs specifically have played a crucial role in sequential modeling tasks and have been used extensively in areas such as natural language processing (NLP), where they frequently outperform their generative counterparts.^14,15^

Recently, deep learning approaches have become popular methods for processing sequential data. However, these models often require a great deal of training data and/or pre-training efforts to show marked improvements over classical ML models.^16^ This is relevant for BGC identification, as the need for experimental characterization of “true” BGCs limits the amount of training data for these approaches; for example, the current version of MIBiG (v2.0) contains only 1,923 experimentally validated BGCs (with some being very closely related to one another, and thus of limited value as training data).^17^ Here, we showed that, with the relatively limited amount of known BGCs available, the linear CRF implemented in GECCO outperforms DeepBGC’s BiLSTM approach, achieving higher accuracy at reduced training and prediction time.

An additional advantage of CRFs over deep learning approaches is that the former are inherently “simpler” and thus more interpretable (whereas “black box” RNNs require substantial additional efforts to “explain” their behavior).^16,26,27^ In the context of BGC mining, an interpretable model can provide insights into genomic mechanisms of secondary metabolism; here, introspection of GECCO’s CRF identified numerous intuitive biological and molecular characteristics that were highly associated with BGC presence. The highly BGC-enriched GO:0042742 and CL0400 terms (corresponding to “defense response to bacterium” and “Double-Glycine leader-peptide cleavage motif”, respectively), for example, are typical of bacteriocin RiPPs often exported by ABC transporters,^28^ while BGC-enriched CL0149 (“CoA-dependent acyltransferase superfamily”) and GO:0008299 (“isoprenoid biosynthetic process”) are associated with polyketide synthases and terpenes, respectively.^29,30^ Furthermore, we identified numerous BGC-associated domains, which had not been included among domain sets previously associated with secondary metabolism, including three highly BGC-associated domains of unknown function. These results not only provide insight into BGC architecture and function, but may be leveraged in the future to improve BGC annotation and identify “high-confidence”, putative novel BGCs, which can be targeted by experimentalists. In conclusion, GECCO’s CRF-based approach used here showcases that model interpretability and computational efficiency can be realised with simultaneous gains in accuracy of *de novo* BGC identification.

## METHODS

### Data acquisition and feature construction

A total of 8,000 randomly selected host-associated prokaryotic contigs were downloaded from the proGenomes2 v12 database^20^ (https://progenomes.embl.de/index.cgi) to serve as candidate BGC-negative instances for training, CV, and testing (accessed 15 July 2020). A Python implementation of the OrthoANI algorithm^31^ (https://github.com/althonos/orthoani) was used to calculate average nucleotide identity (ANI) values between all pairs of candidate contigs. To eliminate the potential risk of training data leakage during CV and testing, a diverse subset of these prokaryotic contigs were selected in which all selected contigs were confirmed to share (i) < 85 ANI with each other and (ii) < 80 ANI with all contigs in the external test set used by Hannigan, et al.^9^ (see section “Validation of CRF performance on external test data” below).

Prodigal v2.6.3^32^ was used to identify ORFs within each of the selected contigs in metagenomic mode (“-p meta”; Supplementary Figure S2). For each contig, the hmmsearch command in HMMER v3.3.1^33^ was used to identify protein domains within the resulting amino acid sequences, using pHMMs from each of the following databases/combinations of databases: (i) Pfam v31.0^34^; (ii) Pfam v32.0^34^; (iii) Pfam v33.1^19^; (iv) TIGRFAM v15.0^18^; (v) PANTHER v15.0^35^; (vi) Pfam v32.0, TIGRFAM v15.0, and PANTHER v15.0; (vii) Pfam v33.1, TIGRFAM v15.0, and PANTHER v15.0; (viii) Pfam v33.1 and TIGRFAM v15.0; (ix) Pfam v33.1, TIGRFAM v15.0, ASPeptides (from antiSMASH v5.1)11, smCOGs (from antiSMASH v5.1),11 and dbCAN v3.0^36^; (x) Pfam v33.1, TIGRFAM v15.0, and Resfams v1.2^37^; (xi) Pfam v33.1, TIGRFAM v15.0, dbCAN v3.0, smCOGs v5.1, and Resfams v1.2; (xii) Pfam v33.1, TIGRFAM v15.0, and smCOGs v5.1; (xiii) Pfam v33.1, TIGRFAM v15.0, smCOGs v5.1, and Resfams v1.2; (xiv) Pfam v33.1 and TIGRFAM v15.1 (Supplementary Figure S2). The resulting ORFs and their respective domains were stored in tabular format and ordered by their start coordinates (referred to hereafter as the “feature table”), and domains with an E-value < 1E-5 were maintained. The command-line implementation of antiSMASH v4.2.0^8^ was then used to identify the coordinates of known BGCs in all selected contigs (using default settings), and ORFs/domains that overlapped with the resulting known BGC regions were removed from the feature table, yielding a final BGC-negative feature table for each prokaryotic contig (Supplementary Figure S2).

To construct a set of BGC-positive instances, the amino acid sequences and metadata for all BGCs within MIBiG v2.0^17^ (https://mibig.secondarymetabolites.org/download) were downloaded (*n =* 1,923). To prevent training data leakage during testing, the diamond blastp command in DIAMOND v0.9.13^38^ was used to align the amino acid sequences of all genomes present in the external test data set (see section “Validation of CRF performance on external test data” below) to the MIBiG BGC amino acid sequences, using minimum amino acid identity (id) and query coverage thresholds (query-cover) of 50% each, and a maximum E-value threshold of 1E-5. MIBiG BGCs were removed from the training set if 50% or more of their amino acid sequences were detected in any test set contigs using DIAMOND and the aforementioned thresholds, yielding a final set of 1,137 MIBiG v2.0 BGCs for training and CV. HMMER was used to identify Pfam domains within the amino acid sequences of the BGCs as described above, producing a BGC-positive feature table for each of 1,137 MIBiG v2.0 BGCs.

To construct a final training set that contained both negative and positive BGC instances, the feature table for a randomly selected MIBiG v2.0 BGC (i.e., a positive instance) was randomly embedded into the feature table of a randomly selected member of the masked, BGC-negative contigs (i.e., a negative instance). This approach yielded a final set of 1,137 contigs that each contained a single MIBiG v2.0 BGC with known coordinates (Supplementary Figure S2).

### CRF training and cross-validation

For each pHMM database combination (*n =* 14; see section “Data acquisition and feature construction” above), a two-state CRF was trained using the CRF architecture implemented in CRFsuite v0.12.^39^ Briefly, for each CRF, features consisted of an ordered list of Python dictionaries, each containing domains identified in each amino acid sequence using the respective pHMMs. Output states corresponded to the probability that a given domain was part of a BGC or not, coded as 1 and 0, respectively (Fig. 1). Additionally, for each pHMM database combination, a feature selection approach was employed, in which the two-sided Fisher’s Exact Test (FET) implemented in the fisher v0.1.9 Python package (implemented as a Cython extension; https://pypi.org/project/fisher) was nested into training fold(s) and used to identify domains associated with BGC presence/absence; the top domains that were associated with the binary outcome variable at a threshold *T* after employing a false-discovery rate correction remained in the model. For each pHMM database combination, values of *T* ranging from 0.05 to 1.0 in increments of 0.05 were tested.

Each combination of pHMM database(s)/feature selection threshold *T* was evaluated using ten-fold CV, using the Kfold function in scikit-learn v0.22.1^40^ and the sequence ID of each ORF treated as a group (i.e., to ensure that each ORF was contained within a single fold and not split across multiple folds; Supplementary Figure S3). For all models that employed it, the FET feature selection approach was nested into training fold(s) to avoid overfitting. Optimization of the *c1* and *c2* CRF hyperparameters (which correspond to L1 and L2 regularization coefficients, respectively) was additionally performed within CV folds, in which either *c1* or *c2* was set to 0.15, while the value of the other hyperparameter was set to one of [0, 0.1, 0.15, 1, 2, 10]. Model performance was evaluated using the following metrics, with scikit-learn and Matplotlib v3.3.4^41^mused to construct all curves: (i) per-protein ROC curves; (ii) per-protein PR curves; (iii) *F*_1_ and (iv) AUPR score versus fraction of FET-selected features. The model selected as the final CRF to be implemented in GECCO (i.e., the CRF trained on BGCs derived from MIBiG v2.0, using domains from Pfam v33.1 and TIGRFAM v15.0, FET inclusion threshold *T =* 0.35, and *c1 = c2 =* 0.15; Supplementary Figure S3) was additionally evaluated using LOTO CV for each MIBiG biosynthetic class, with BGCs assigned to multiple biosynthetic classes excluded (Supplementary Figure S8).

### Validation of CRF performance on external test data

The (i) six genome test set and (ii) nine genome bootstrap set used by Hannigan, et al.^9^ (see Supplementary Tables S4 and S3 of Hannigan, et al.^9^, respectively) were used as external test sets to evaluate the performance of the GECCO CRF (see section “CRF training and cross-validation” above). To construct an extensive third external test set comprising known BGC and non-BGC regions, BGCs that were present in MIBiG v2.0 but absent from MIBiG v1.3 were each embedded into a randomly selected prokaryotic contig as described above (see section “Data acquisition and feature construction” above; Supplementary Figure S2). For this external test set, DIAMOND was used to identify potentially redundant BGCs in MIBiG v2.0 that aligned to BGCs in MIBiG v1.3, using the blastp thresholds described above (see section “Data acquisition and feature construction” above); contigs that contained these potentially redundant BGCs were removed from the external test set to avoid training data leakage during testing, yielding a final set of 376 contigs that each contained a single BGC present in MIBiG v2.0 but absent from MIBiG v1.3 (referred to as the “376-genome test set”).

To avoid training leakage into the 376-genome test set, the GECCO CRF (see section “CRF training and cross-validation” above) was re-trained on BGCs available in MIBiG v1.3 and was used to predict BGC presence/absence in each genome in the three test sets (i.e., the six-, nine-, and 376-genome test sets; Fig. 2 and Supplementary Figures S4-S7). The ability of each of the following methods to predict BGC presence/absence was additionally evaluated on each of the three test sets (Fig. 2a-c,e and Supplementary Figure S6): (i) DeepBGC v0.1.18^9^; (ii) the original ClusterFinder^7^ algorithm, implemented in DeepBGC v0.1.18; (iii) the retrained version of the ClusterFinder algorithm, implemented in DeepBGC v0.1.18 (re-trained on BGCs available in MIBiG v1.3); (iv) a re-trained implementation of DeepBGC, which was trained on the exact positive and negative BGC instances used to retrain the GECCO CRF, using BGCs available in MIBiG v1.3 (using DeepBGC’s “train” function). The re-trained implementation of DeepBGC (iv) was additionally evaluated relative to the re-trained implementation of the GECCO CRF (i.e., trained on BGCs from MIBiG v1.3) using 10-fold CV, where both models were trained and tested on identical folds (see section “CRF training and cross-validation” above; Supplementary Figure S5). For all models, performance was evaluated using: per-domain (i) ROC and (ii) PR curves; (iii) segment overlap PR curves (Supplementary Figure S4), using minimum overlap thresholds of 25, 50, and 75% (Fig. 2a-c, Supplementary Figure S6-S7).

### Prediction of biosynthetic class

To assign the BGCs that the GECCO CRF predicted to one or more of the six biosynthetic classes in MIBiG v2.0 (with MIBiG’s “Other” class excluded), the following classifiers were trained (Supplementary Figure S9): (i) a random forest classifier, using the scikit-learn RandomForestClassifier function; (ii) an ExtraTrees classifier, using the scikit-learn ExtraTreesClassifier function; (iii) a *k*-nearest neighbors (kNN) classifier, using the scikit-learn KNeighborsClassifier function, a cosine distance metric, and number of neighbors *n =* 3; (iv) the aforementioned kNN, with *n =* 15. For each classifier, BGCs were represented by compositional vectors, where individual features corresponded to the fraction of a particular domain present in the BGC. For example, a predicted BGC with 2 domains *A*, one domain *B*, and one domain *C* would be represented by domain composition vector [*A*: 0.5, *B*: 0.25, *C*: 0.25], assuming *A, B,* and *C* are the only possible domains. The ability of each classifier to predict MIBiG biosynthetic class was evaluated using five-fold CV via the cross_val_predict function in scikit-learn, and the random forest was implemented as the final biosynthetic classifier in GECCO (Supplementary Figure S9).

### BGC identification in prokaryotic genomes

Each of the following methods was used to identify BGCs in all representative genomes available in the proGenomes2 v12 database^20^ (*n* = 12,221; accessed 15 July 2020): (i) the GECCO CRF trained on BGCs available in MIBiG v2.0 (i.e., the final model implemented in GECCO, run using default parameters); (ii) antiSMASH v4.2.0^8^ (run using default parameters); (iii) DeepBGC v0.1.18^9^ (run using default parameters with the addition of DeepBGC’s “--prodigal-meta-mode” option, as GECCO uses this option for BGC detection by default; Fig. 3ab). antiSMASH-to-MIBiG type mappings from BiG-SLiCE v1.1.0^21^ were used to map antiSMASH biosynthetic types to MIBiG biosynthetic types (used by DeepBGC and GECCO; Supplementary Table S2). The three aforementioned BGC detection/classification methods were additionally applied to the following data sets to assess their speed using a single CPU (Fig. 2f and Supplementary Figure S10): (i) contigs in each of the three test sets (i.e., the six-, nine-, and 376-genome test sets, *n =* 395 contigs; see section “Validation of CRF performance on external test data” above); (ii) the 12,221 proGenomes2 representative genomes. Plots were constructed in R v3.6.1^42^ using ggplot2 v3.3.3.^43^

### Comparison of GECCO and BiG-SLiCE domain sets

Domains that were included in the optimized GECCO pHMMs based on their FET association with BGC presence/absence (see section “CRF training and cross-validation” above) were compared to protein domains used by BiG-SLiCE v1.1.0.^21^ BiG-SLiCE, which is designed to cluster antiSMASH BGCs into Gene Cluster Families, relies on a set of core biosynthetic domains for BGC annotation and clustering. Domains within the GECCO pHMMs were compared to all publicly available core biosynthetic BiG-SLiCE domains with a reported accession number, as well as BiG-SLiCE’s larger set of “BioPfam” domains (identical to the Pfam v33.1 database) by finding the union of the three domain sets and plotting via venn.js (https://github.com/benfred/venn.js/) and Matplotlib (Fig. 3c and Supplementary Table S1). Three independent, two-group Mann-Whitney *U* tests were used to compare CRF weights associated with the following GECCO domain sets, using the “wilcox.test” function in R, with parameters set to perform an unpaired (paired = F), two-sided (alternative = “two.sided“) test using a normal approximation (exact = F) and a continuity correction (correct = T): (i) GECCO domains included in BiG-SLiCE’s core biosynthetic domain set; (ii) GECCO Pfam domains excluded from BiG-SLiCE’s core biosynthetic domain set; (iii) GECCO Tigrfam domains excluded from BiG-SLiCE’s core biosynthetic domain set. Tests between groups (i)/(iii) and (ii)/(iii) were statistically significant after a Bonferroni correction (raw *P =* 3.12e-07 and 5.75e-06, respectively), but not groups (i)/(ii) (raw *P =* 0.10).

### GO term enrichment

Weights associated with each protein domain were extracted from the trained GECCO CRF instance, and all available GO terms for each domain were retrieved from InterPro (*n =* 2,722 domains with one or more assigned GO terms, out of 5,255 total domains).^22,44^ To identify over-represented GO terms associated with BGC presence (i.e., BGC-enriched GO terms), domains were assigned ranks based on their weights, where the domain with the highest weight (i.e., PF14867, with weight 4.190953) was assigned a value of “1”, and the domain with the lowest weight (i.e., PF02881, with weight -1.798162) was assigned a value of “2722”. For each of the (i) Biological Process and (ii) Molecular Function GO ontologies, the runTest function in the topGO v2.36.0 package^45^ in R v3.6.1 was used to perform a Kolmogorov-Smirnov (KS) test (statistic = “ks”), using the “weight01” algorithm (algorithm = “weight01”) to account for the GO graph topology.^45^ Enrichment scores were calculated for all statistically significant (*P <* 0.05) GO terms by negating the base-10 logarithm of the resulting *P*-values. The aforementioned steps were repeated to identify over-represented GO terms associated with BGC-absence, using (i) domains ranked by weight from lowest-to-highest (i.e., the domain with the lowest weight was assigned a value of “1”, and the domain with the highest weight was assigned a value of “2722”) and (ii) enrichment scores corresponding to the non-negated base-10 logarithms of the resulting *P*-values. topGO’s weight01 algorithm calculates the *P*-value of a GO term conditioned on neighbouring GO terms; therefore, tests were considered not independent, and *P*-values were interpreted as inherently corrected.^45^ Enrichment scores were plotted using the ggplot2 package in R (Fig. 3d and Supplementary Figure S11).

### Pfam clan enrichment

Weights associated with each Pfam protein domain were extracted from the trained GECCO CRF instance, and all available Pfam domain-to-clan mappings were retrieved for Pfam v33.1 via FTP (*n =* 1,907 Pfam domains with an assigned clan, out of 2,873 total Pfam domains).^19^ A (i) vector of raw Pfam domain weights (ordered from highest-to-lowest) and (ii) list of clan-to-domain mappings were supplied to the fgsea function from the fgsea v1.10.1 R package^46,47^, which was used to identify BGC- and non-BGC-enriched Pfam clans, using 1 million permutations (nperm = 1000000), a minimum clan size of three (minSize = 3), and no maximum clan size limit. For significantly enriched clans (false discovery rate-corrected *P <* 0.10), ggplot2 was used to plot (i) fgsea normalized enrichment scores (NES) and (ii) the negated base-10 logarithm of the false discovery rate-corrected *P*-values (Fig. 3e and Supplementary Figure S12)

## Data availability

Training and test data can be downloaded from https://github.com/zellerlab/GECCO/releases/tag/v0.6.0. GECCO CRF weights are available in Supplementary Table S1.

### Code availability

GECCO code is free and publicly available at https://gecco.embl.de.

## ACKNOWLEDGMENTS

We are grateful to Tobias Gulder, Maximilian Hohmann, and members of the Zeller Team for fruitful discussions, as well as EMBL IT Services for support with high-performance computing. This work was funded by the European Molecular Biology Laboratory, the German Research Foundation (Deutsche Forschungsgemeinschaft, DFG, grant no. 395357507 – SFB 1371), and the German Federal Ministry of Education and Research (BMBF, grant no. 031L0181A, and the de.NBI network, grant no. 031A537B).

## AUTHOR CONTRIBUTIONS

Software development and computational analyses were performed by JSF, ML, and LMC with contributions of data or tools from all authors. GZ conceived and funded the study. LMC and GZ co-wrote the manuscript with input from all authors.

