## Supplementary Figure S1 for "Accurate *de novo* identification of biosynthetic gene clusters with GECCO"

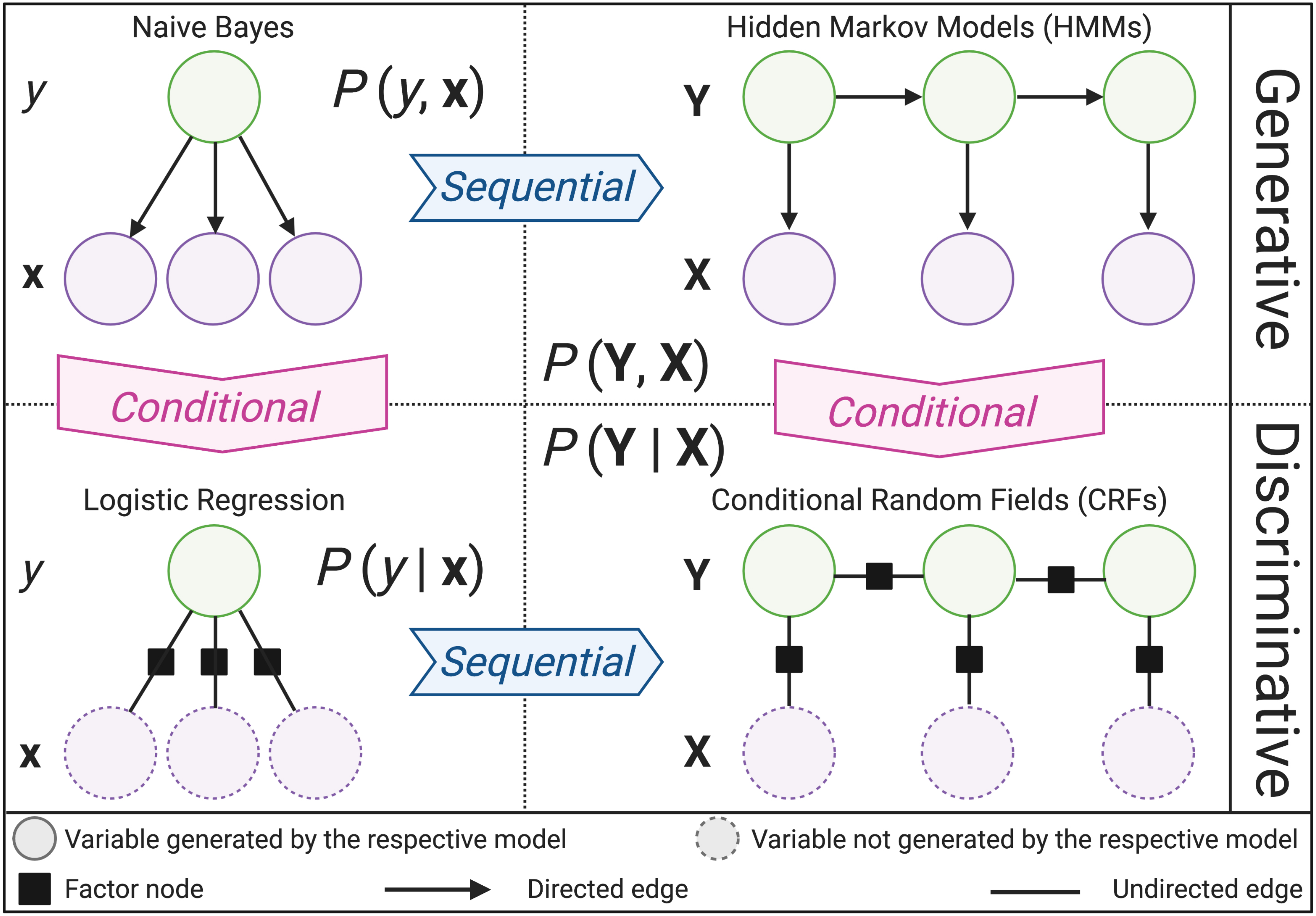


**Supplementary Figure S1.** High-level comparison of the graphical structures of Naïve Bayes, logistic regression, linear hidden Markov models (HMMs), and linear conditional random fields (CRFs). The figure was adapted from Figure 2.3 of Sutton and McCallum, 2010 (Sutton, C. and McCallum, A., 2010, "An Introduction to Conditional Random Fields for Relational Learning", *arXiv:1011.4088v1*). The figure was created with BioRender.com.
