## Supplementary Figure S2 for "Accurate *de novo* identification of biosynthetic gene clusters with GECCO"

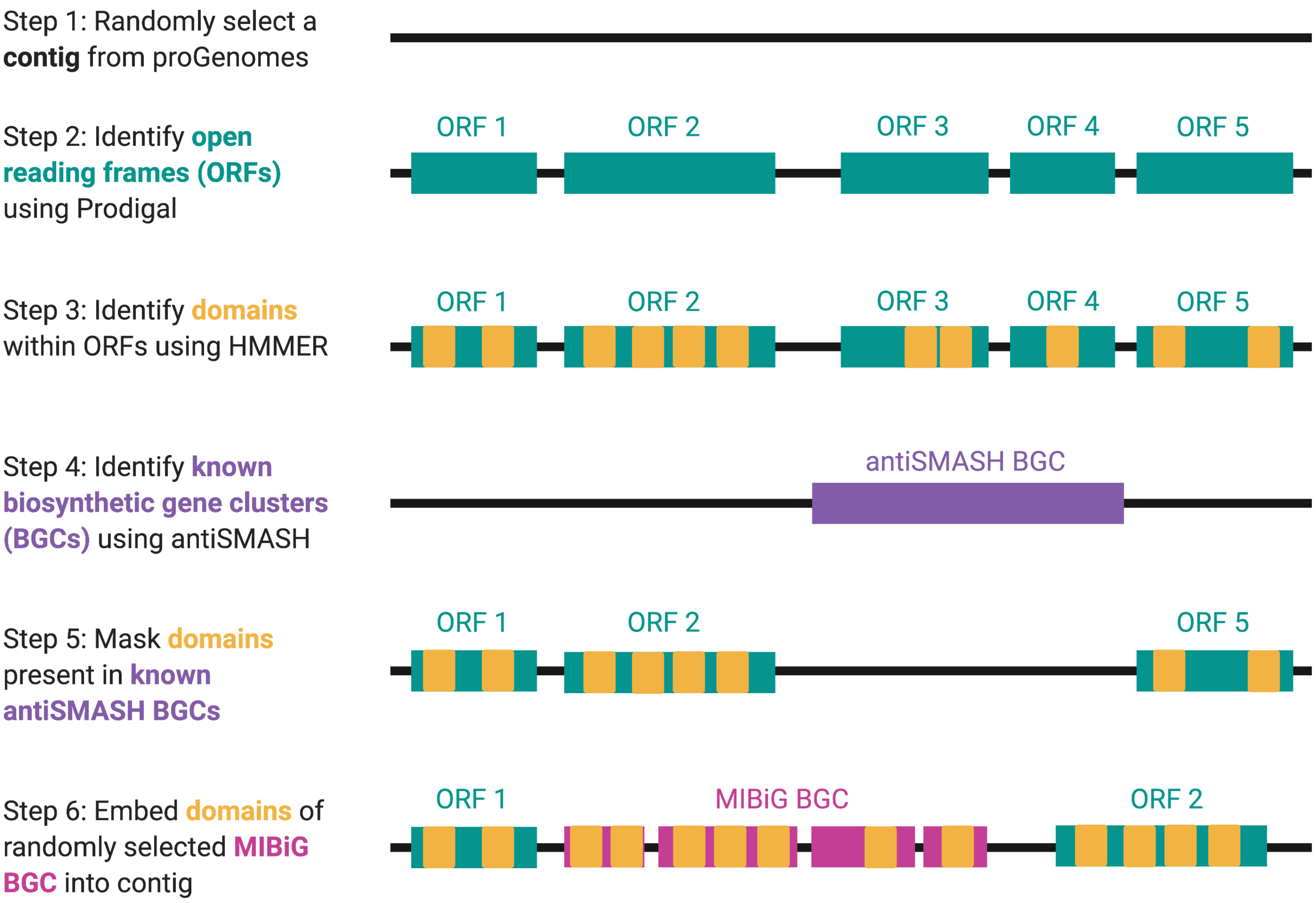


**Supplementary Figure S2.** Illustrated overview of the methodology used to construct the training data sets for GECCO and the re-trained implementation of DeepBGC described in the manuscript. Briefly, open reading frames (ORFs) were identified in a randomly selected bacterial contig (Steps 1 and 2). Protein domains were identified within the resulting set of ORFs using HMMER and profile hidden Markov models (pHMMs) from one or more protein domain databases (Step 3). The same bacterial contig was supplied to antiSMASH, which was then used to identify known BGCs using its rule-based BGC detection approach (Step 4). Protein domains detected within antiSMASH BGC regions were removed, yielding a set of ordered domains with no known BGCs (i.e., BGC-negative training instances; Step 5). A BGC present in the Minimum Information about a Biosynthetic Gene cluster (MIBiG) database was then randomly selected and randomly embedded into the bacterial contig (Step 6), yielding a set of contigs with one known MIBiG BGC per contig. The figure was created with BioRender.com.
