## Supplementary Figure S3 for "Accurate *de novo* identification of biosynthetic gene clusters with GECCO"

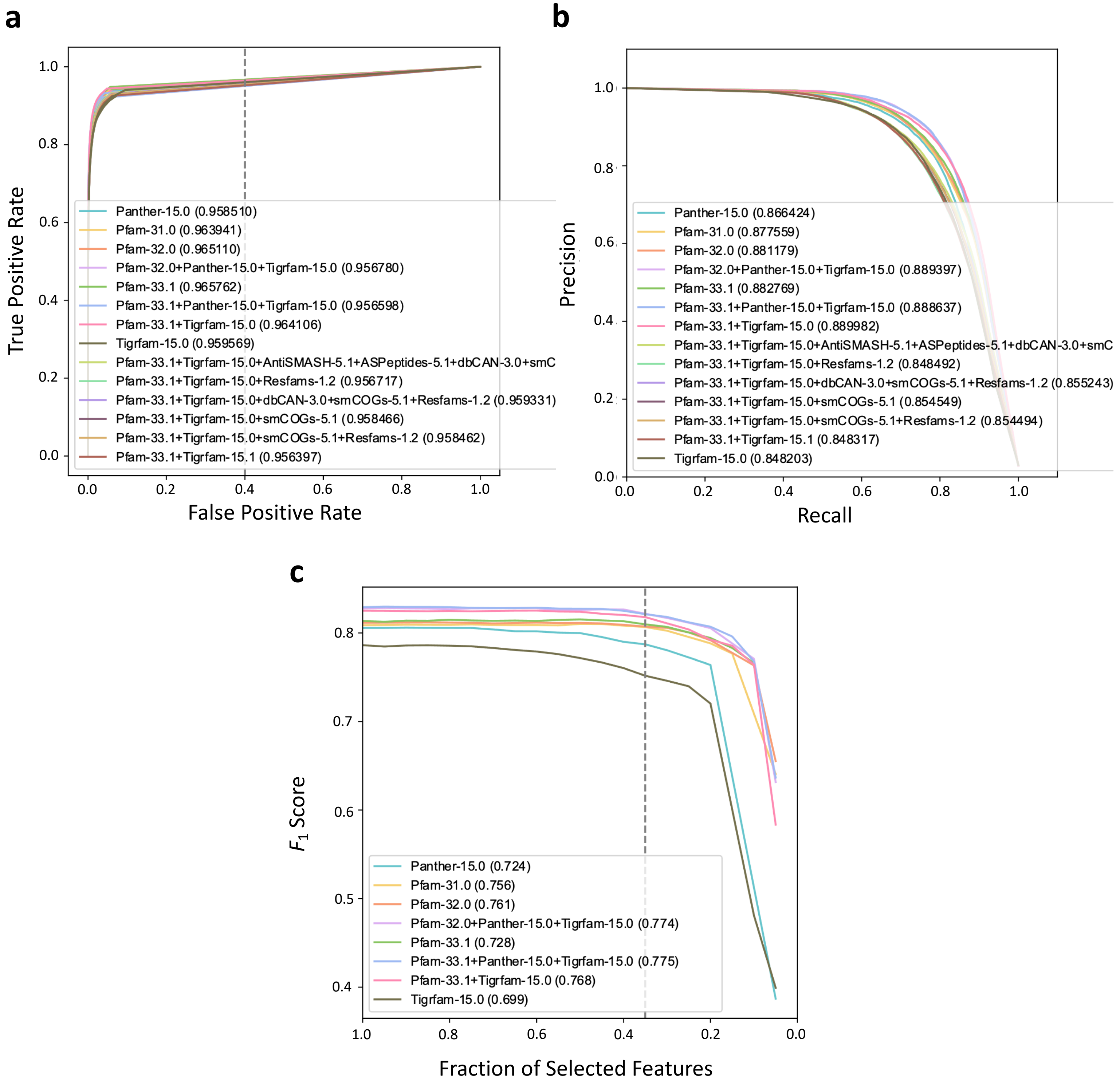


**Supplementary Figure S3.** (a) Protein-level receiver operating characteristic (ROC) and (b) protein-level precision-recall (PR) curves constructed using results of 10-fold cross validation (CV) for conditional random fields (CRFs) trained on features derived from one or more of the following protein domain databases: (i) Pfam v31.0 (Pfam-31.0); (ii) Pfam v32.0 (Pfam-32.0); (iii) Pfam v33.1 (Pfam-33.1); (iv) TIGRFAM v15.0 (Tigrfam-15.0); (v) PANTHER v15.0 (Panther-15.0); (vi) antiSMASH v5.1 (AntiSMASH-5.1); (vii) ASPeptides (from antiSMASH v5.1; ASPeptides-5.1); (viii) smCOGs (from antiSMASH v5.1; smC/smCOGs-5.1); (ix) Resfams v1.2 (Resfams-1.2); (x) dbCAN v3.0 (dbCAN-3.0). Area under the curve (AUC) values associated with each model are reported in the legend, next to the name of the respective model. (c) Protein-level curves showcasing F1 score (Y-axis) versus the ratio of Fisher's Exact Test (FET)-selected features (X-axis) included in CRFs trained on protein domains from the aforementioned databases. The final FET feature selection threshold chosen for the optimal GECCO model is denoted by the vertical gray dashed line (i.e., FET inclusion threshold of 0.35). AUC values associated with each model are reported in the legend, next to the name of the respective model. 10-fold CV was performed on the BGCs available in MIBiG v2.0, each of which were embedded into a randomly selected bacterial contig. The model chosen as the final CRF model implemented in GECCO is the "Pfam-33.1+Tigrfam-15.0" model with a 0.35 FET inclusion threshold.
