## Supplementary Figure S4 for "Accurate *de novo* identification of biosynthetic gene clusters with GECCO"

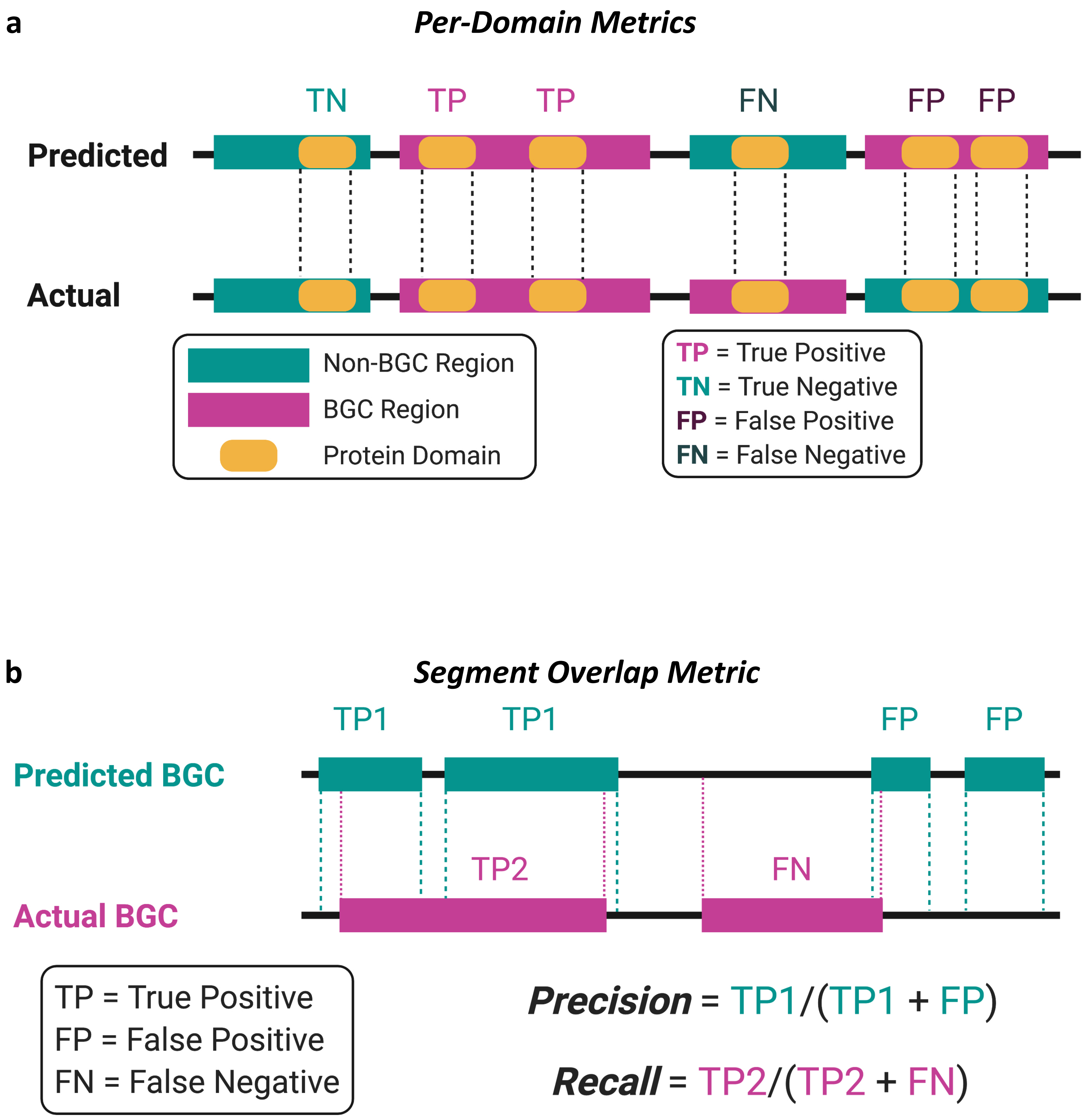


**Supplementary Figure S4.** (a) Graphical depiction of the methodology used to calculate per-domain metrics ($F_1$ scores, per-domain receiver operating characteristic [ROC] curves, per-domain precision-recall [PR] curves). (b) Graphical depiction of the methodology used to calculate segment overlap PR metrics. Predicted and actual biosynthetic gene clusters (BGCs) detected within a contig (solid black lines) are denoted by teal and magenta boxes, respectively. Dashed lines denote BGC boundaries. BGCs that overlapped by a given overlap threshold or more were considered to be "overlapping BGCs". Overlap thresholds of 25, 50, and 75% were tested. The figure was created with BioRender.com.
