## Supplementary Figure S5 for "Accurate *de novo* identification of biosynthetic gene clusters with GECCO"

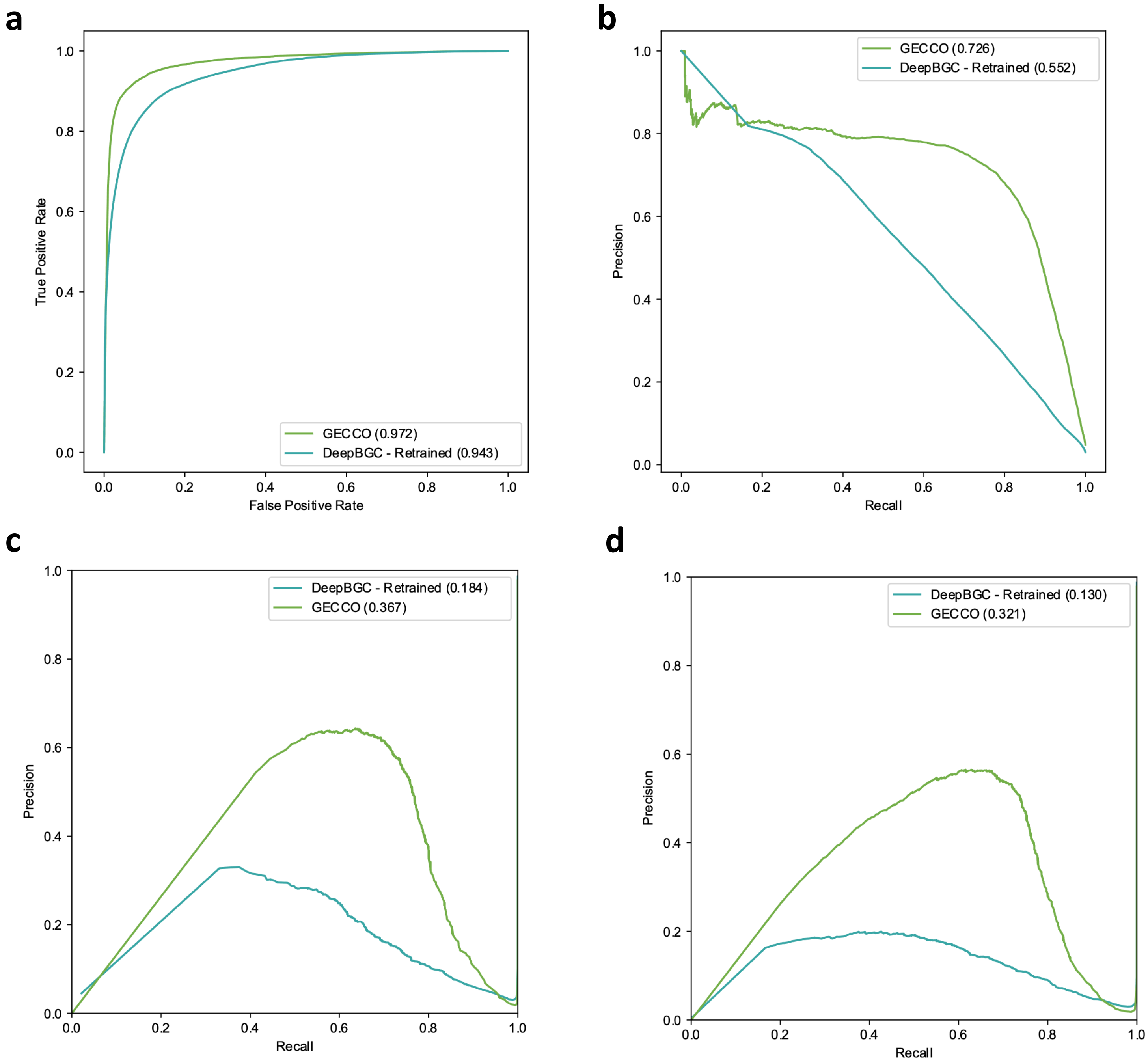


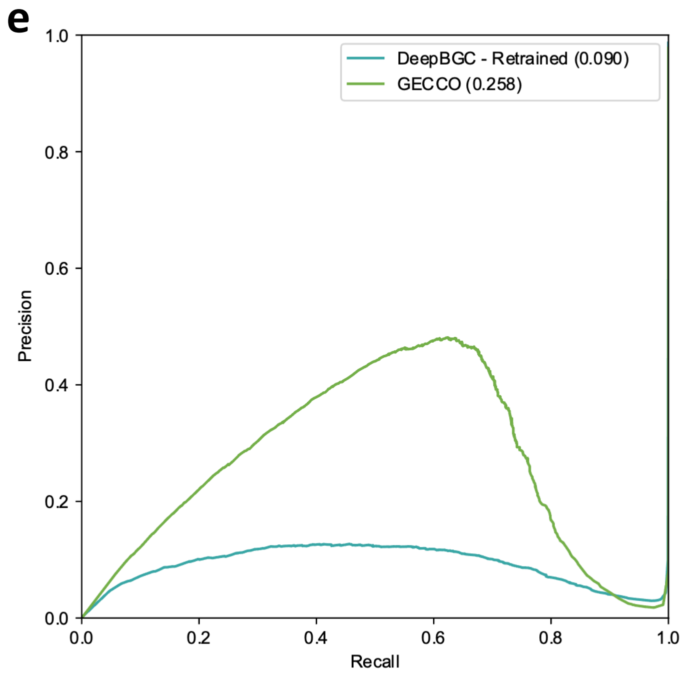


**Supplementary Figure S5.** Results of 10-fold cross validation (CV), showcased using (a) domain-level receiver operating characteristic (ROC) and (b) domain-level precision-recall (PR) curves, as well as segment-overlap PR curves constructed using minimum overlap thresholds of (c) 25, (d) 50, and (e) 75% overlap (see Supplementary Figure S4 for details regarding how the segment overlap values were calculated). All subplots (a-e) were constructed using results of 10-fold CV for each of the following methods: (i) a retrained implementation of DeepBGC (DeepBGC - Retrained); (ii) GECCO, a CRF trained using positive BGC instances derived from MIBiG v1.3, an optimized subset of domains from Pfam v33.1 and Tigrfam v15.0, and a Fisher's Exact Test (FET) feature inclusion threshold ($T$) of 35% (i.e., $T = 0.35$). Both methods were trained and evaluated on identical data/folds. Area under the curve (AUC) values associated with each model are reported in the legend, next to the name of the respective model.
