## Supplementary Figure S6 for "Accurate *de novo* identification of biosynthetic gene clusters with GECCO"

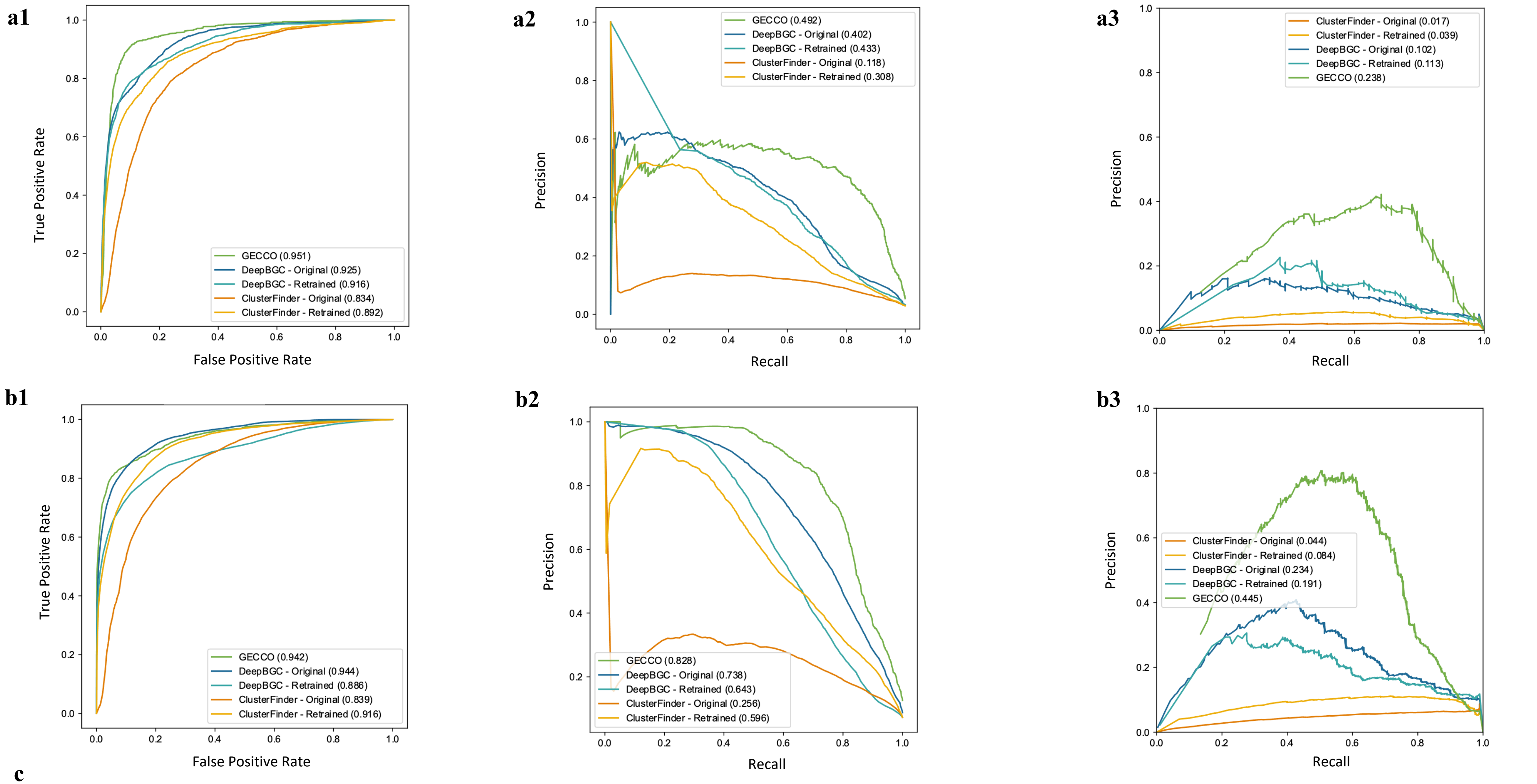


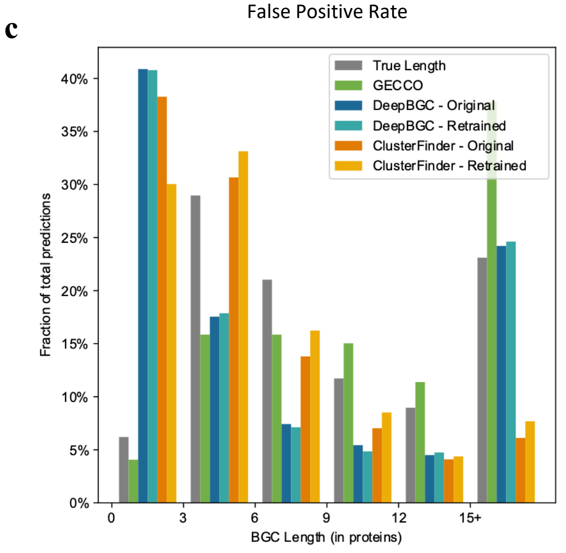


**Supplementary Figure S6.** (1) Domain-level receiver operating characteristic (ROC), (2) domain-level precision recall (PR), and (3) segment-overlap PR curves constructed using biosynthetic gene clusters (BGCs) detected in the (a) six- and (b) nine-genome test sets for each of the original and retrained implementations of (i) ClusterFinder (ClusterFinder-Original and ClusterFinder-Retrained, respectively) and (ii) DeepBGC (DeepBGC-Original and DeepBGC-Retrained, respectively). "GECCO" denotes a CRF trained on MIBiG v1.3, using an optimized subset of Pfam v33.1 and Tigrfam v15.0 domains and a Fisher's Exact Test (FET) feature inclusion threshold ($T$) of 35% (i.e., $T = 0.35$). Segment-overlap PR curves (3) were constructed using 50% segment overlap (see Supplementary Figure S4 for details regarding how the segment overlap values were calculated). Area under the curve (AUC) values associated with each model are reported in the legend. (c) Histogram of predicted BGC lengths (in number of proteins; X-axis) relative to "true" BGC lengths (True Length) among genomes in the six- and nine-genome test sets. The Y-axis denotes the percentage of total BGC predictions.
