## Supplementary Figure S7 for "Accurate *de novo* identification of biosynthetic gene clusters with GECCO"

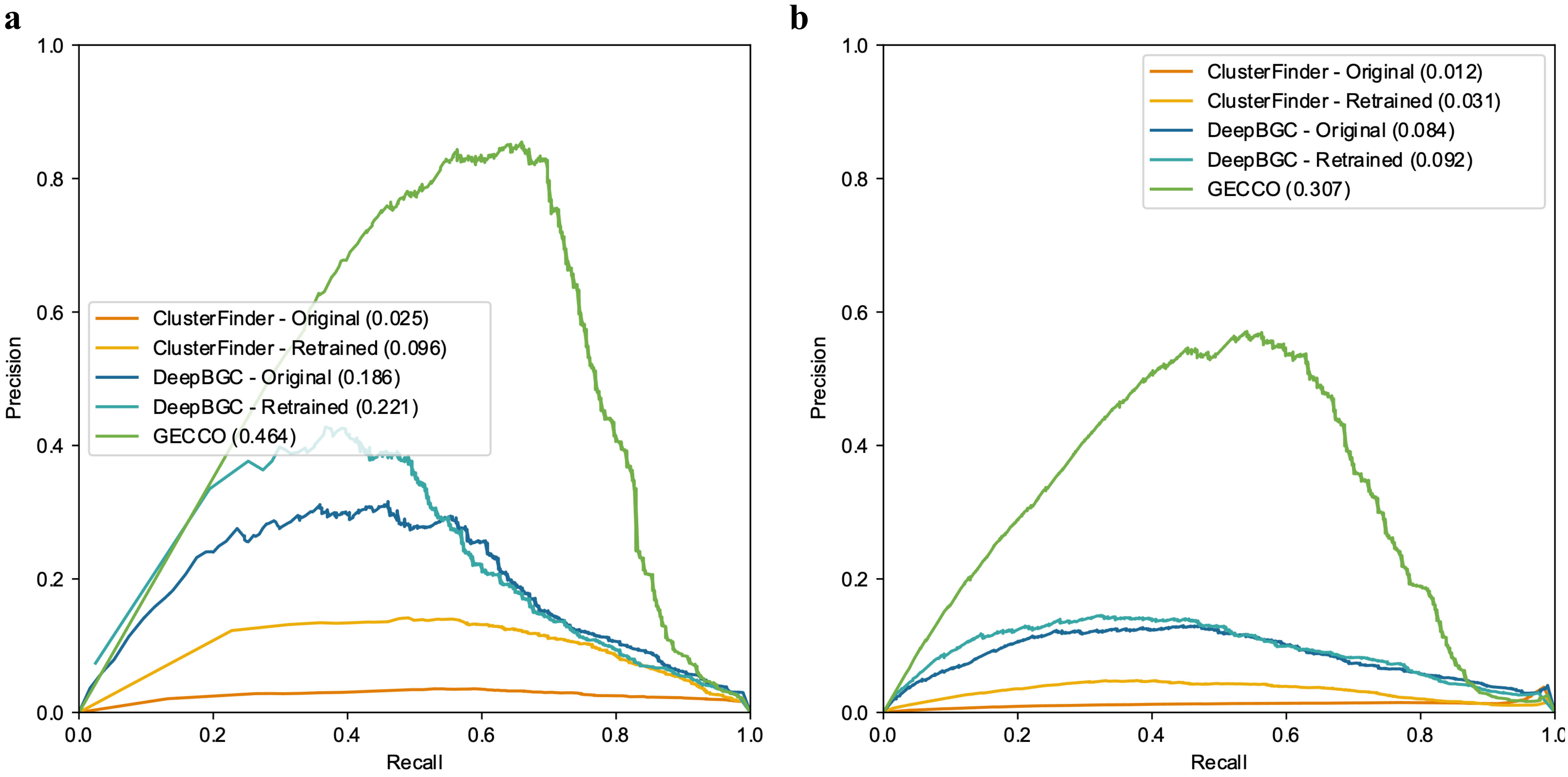


**Supplementary Figure S7.** Segment-overlap precision-recall (PR) curves constructed using biosynthetic gene clusters (BGCs) detected in the 376-genome test set for original and retrained implementations of (i) ClusterFinder (ClusterFinder-Original and ClusterFinder-Retrained, respectively) and (ii) DeepBGC (DeepBGC-Original and DeepBGC-Retrained, respectively). "GECCO" denotes a CRF trained on a subset of Pfam v33.1 and Tigrfam v15.0 domains, selected using a Fisher's Exact Test (FET) feature inclusion threshold ($T$) of 35% (i.e., $T = 0.35$). Segment-overlap PR curves were constructed using (a) 25 and (b) 75% overlap (see Supplementary Figure S4 for details regarding how the segment overlap values were calculated). All models (ab) were trained on BGCs from MIBiG v1.3, and area under the curve (AUC) values are reported in legends.
