## Supplementary Figure S8 for "Accurate *de novo* identification of biosynthetic gene clusters with GECCO"

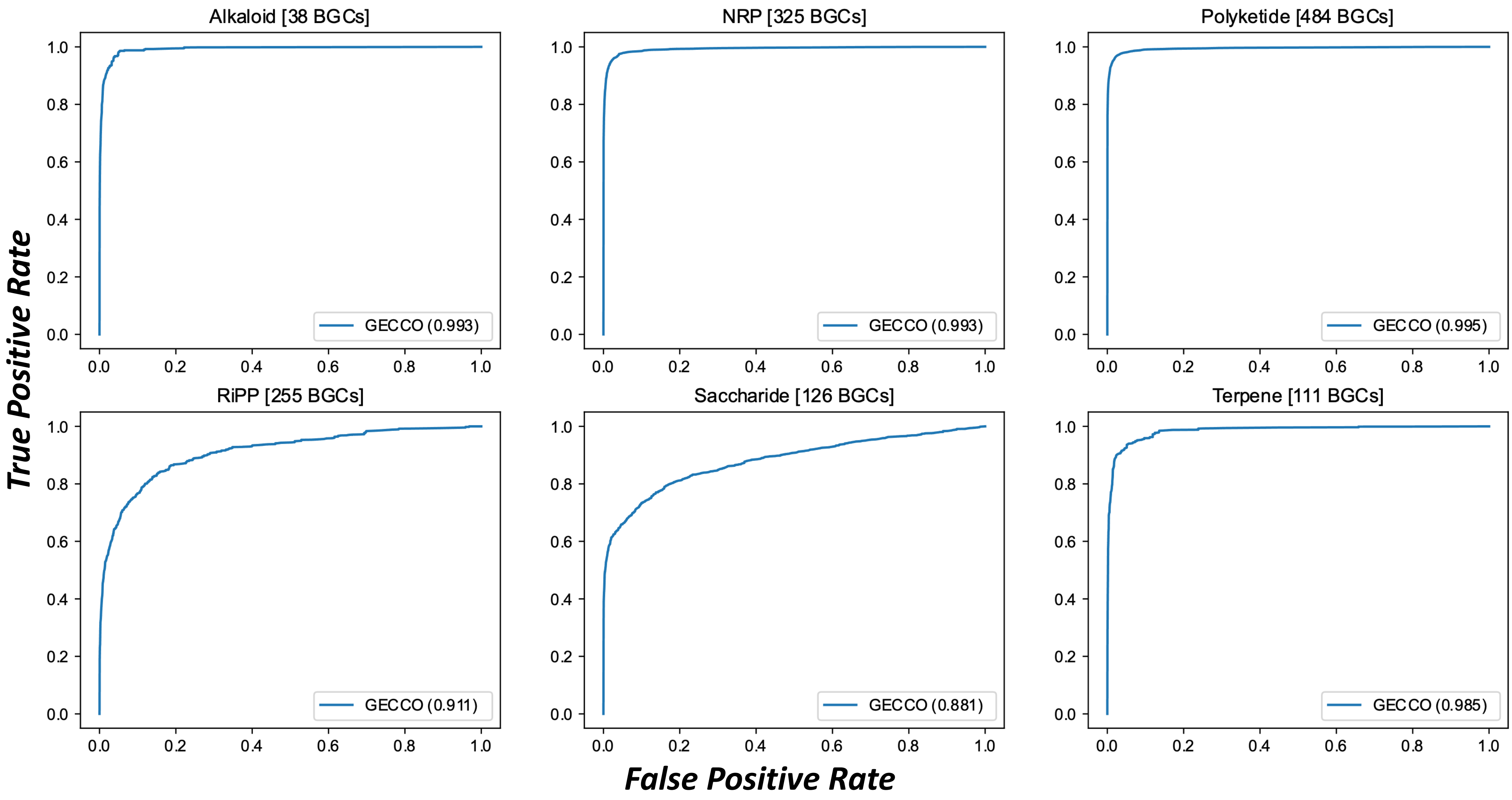


**Supplementary Figure S8.** Receiver operating characteristic (ROC) curves constructed using the results of leave-one-type-out (LOTO) cross-validation (CV) for the optimal GECCO conditional random field (CRF; i.e., features derived from a subset of protein domains from the Pfam v33.1 and TIGRFAM v15.0 databases, using a Fisher's Exact Test inclusion threshold of 0.35). Biosynthetic gene clusters (BGCs) were derived from MIBiG v2.0 and were embedded in a randomly selected bacterial contig (Supplementary Figure S2). LOTO CV "types" correspond to MIBiG biosynthetic classes and are included in the sublot titles. Area under the curve (AUC) values associated with each LOTO CV MIBiG biosynthetic class are reported in each subplot legend. MIBiG's "Other" class was not included in the evaluation. NRP, nonribosomal peptide; RiPP, ribosomally synthesized and post-translationally modified peptide.
