## Supplementary Figure S9 for "Accurate *de novo* identification of biosynthetic gene clusters with GECCO"

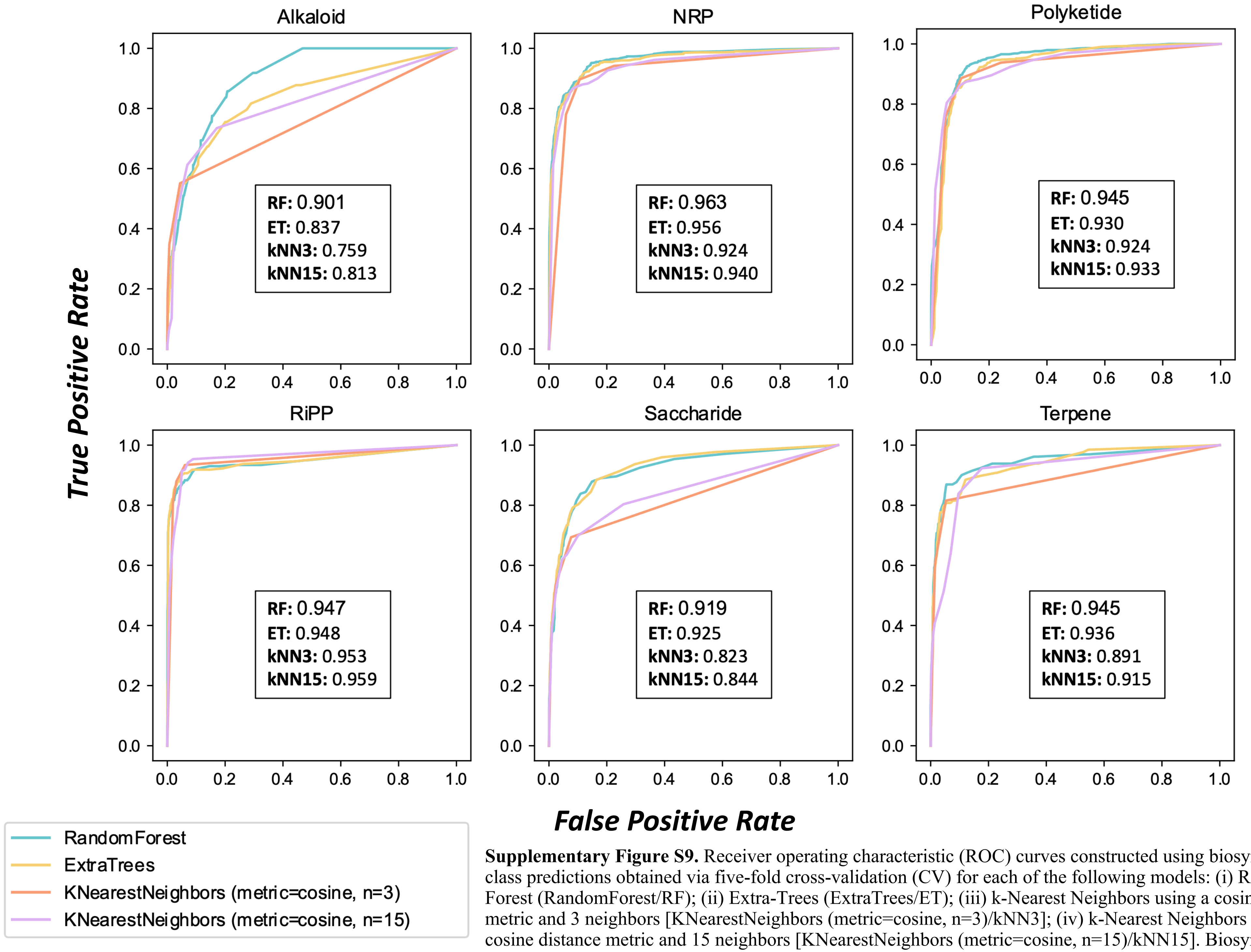


**Supplementary Figure S9.** Receiver operating characteristic (ROC) curves constructed using biosynthetic class predictions obtained via five-fold cross-validation (CV) for each of the following models: (i) Random Forest (RandomForest/RF); (ii) Extra-Trees (ExtraTrees/ET); (iii) k-Nearest Neighbors using a cosine distance metric and 3 neighbors [KNearestNeighbors (metric=cosine, n=3)/kNN3]; (iv) k-Nearest Neighbors using a cosine distance metric and 15 neighbors [KNearestNeighbors (metric=cosine, n=15)/kNN15]. Biosynthetic gene clusters (BGCs) were derived from MIBiG v2.0 and were each embedded into a randomly selected bacterial contig (Supplementary Figure S2). Biosynthetic classes correspond to MIBiG biosynthetic classes and are included in the sublot titles. Area under the curve (AUC) values associated with each model are reported in the legend. NRP, nonribosomal peptide; RiPP, ribosomally synthesized and post-translationally modified peptide.
