## Supplementary Figure S10 for "Accurate *de novo* identification of biosynthetic gene clusters with GECCO"

a

b

c

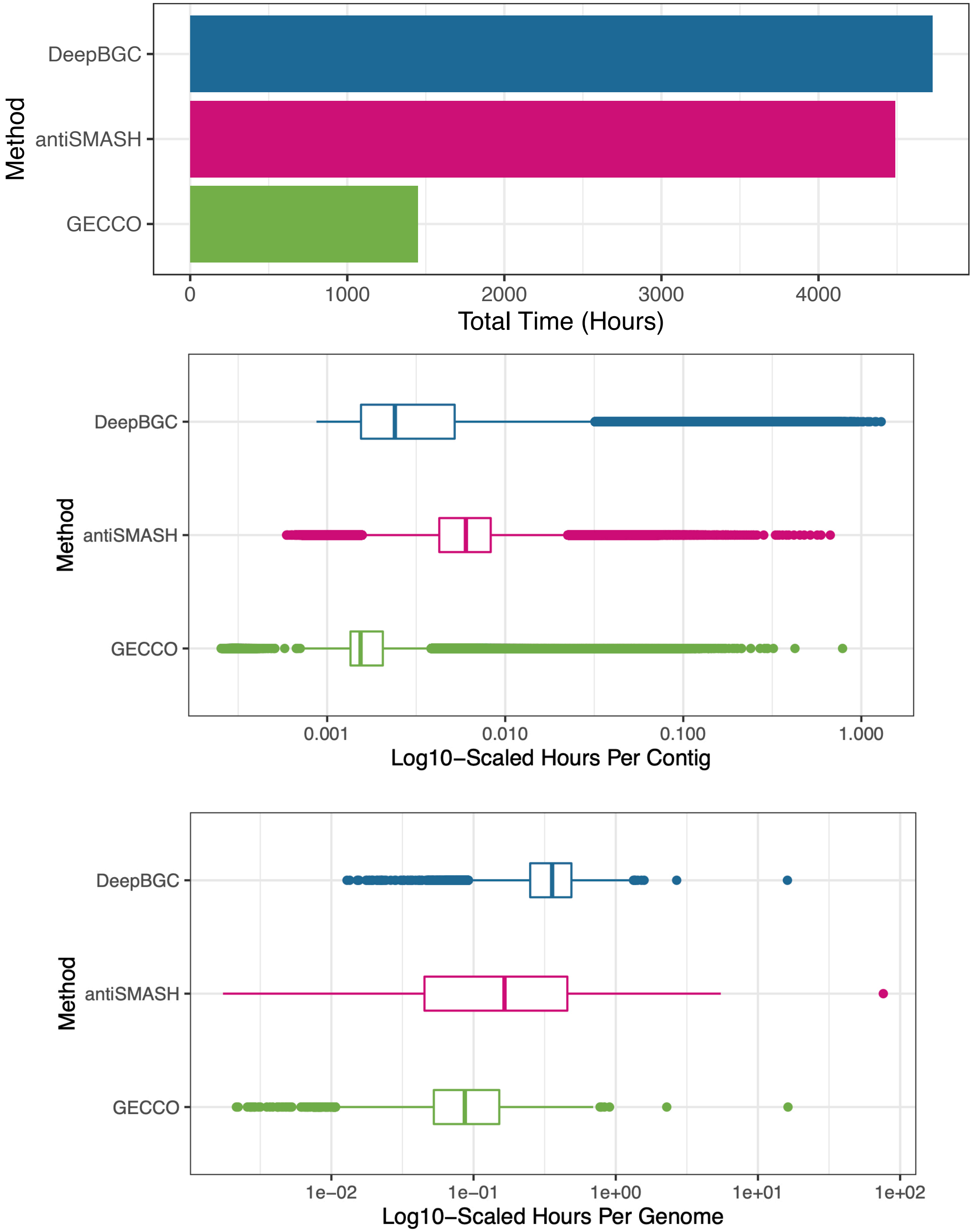


**Supplementary Figure S10.** (a) Runtime required to detect and classify BGCs in all proGenomes2 representative genomes (*n* = 12,221) with one CPU, using antiSMASH, DeepBGC, and GECCO. (bc) Base-10 logarithm-scaled time in hours (X-axis) required to identify and classify biosynthetic gene clusters (BGCs) in all representative genomes from the proGenomes2 database using GECCO, DeepBGC, and antiSMASH, and one CPU. Times are reported per (b) contig (*n* = 627,182 contigs) and (c) genome (*n* = 12,221 genomes). For all plots, antiSMASH v5.1.211, DeepBGC v0.1.18, and GECCO v0.6.0 were used.
