## Supplementary Figure S11 for "Accurate *de novo* identification of biosynthetic gene clusters with GECCO"

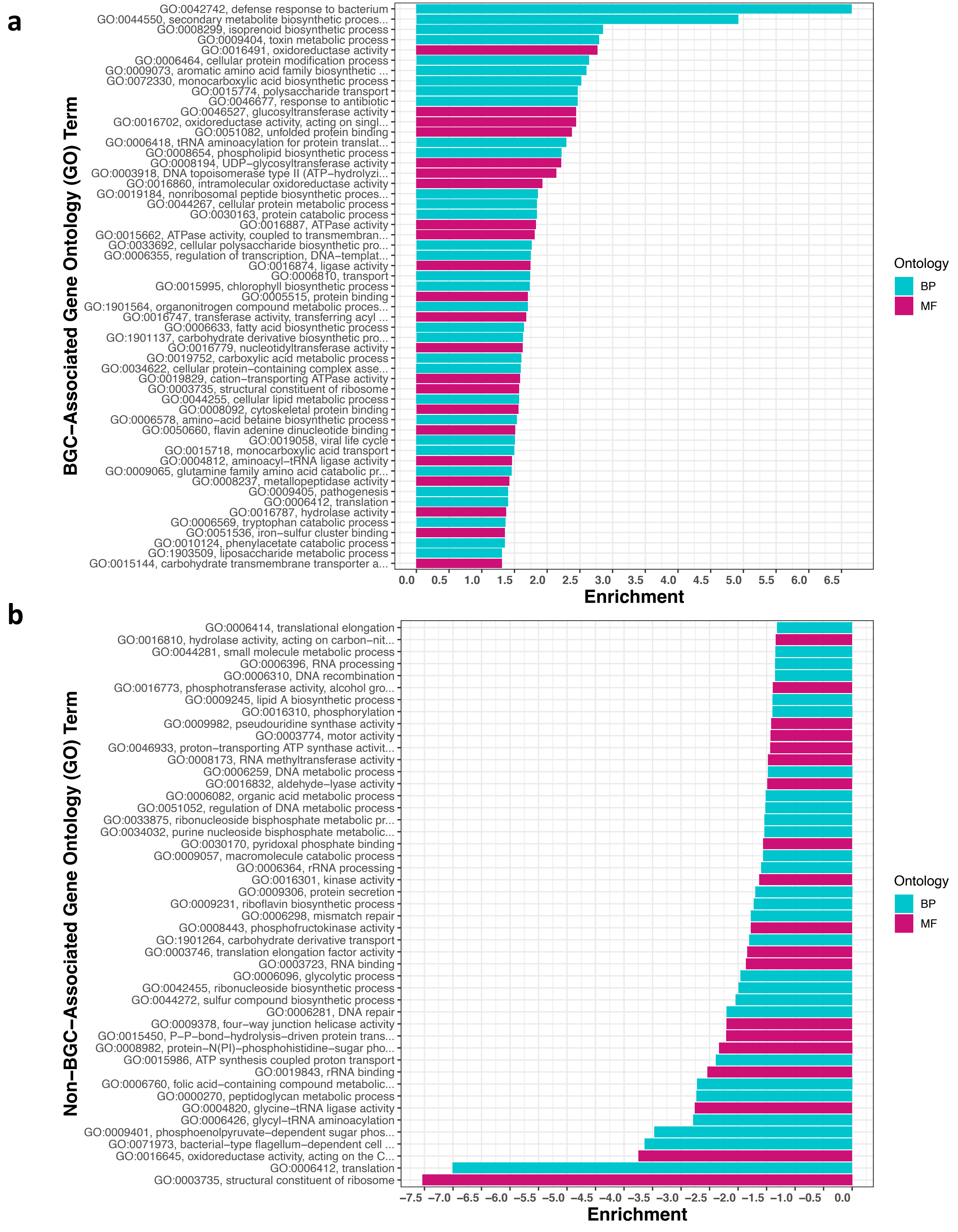


**Supplementary Figure S11.** Gene Ontology (GO) terms (Y-axis) enriched in (a) BGC and (b) non-BGC regions, obtained using the Kolmogorov-Smirnov test/weight01 algorithm implemented in topGO ($P < 0.05$). Enrichment scores (X-axis) were calculated by taking the (a) negated and (b) non-negated base-10 logarithm of each GO term's *P*-value. Bars are colored by the ontology from which the respective GO term was derived. BP, Biological Process; MF, Molecular Function.
