## Supplementary Figure S12 for "Accurate *de novo* identification of biosynthetic gene clusters with GECCO"

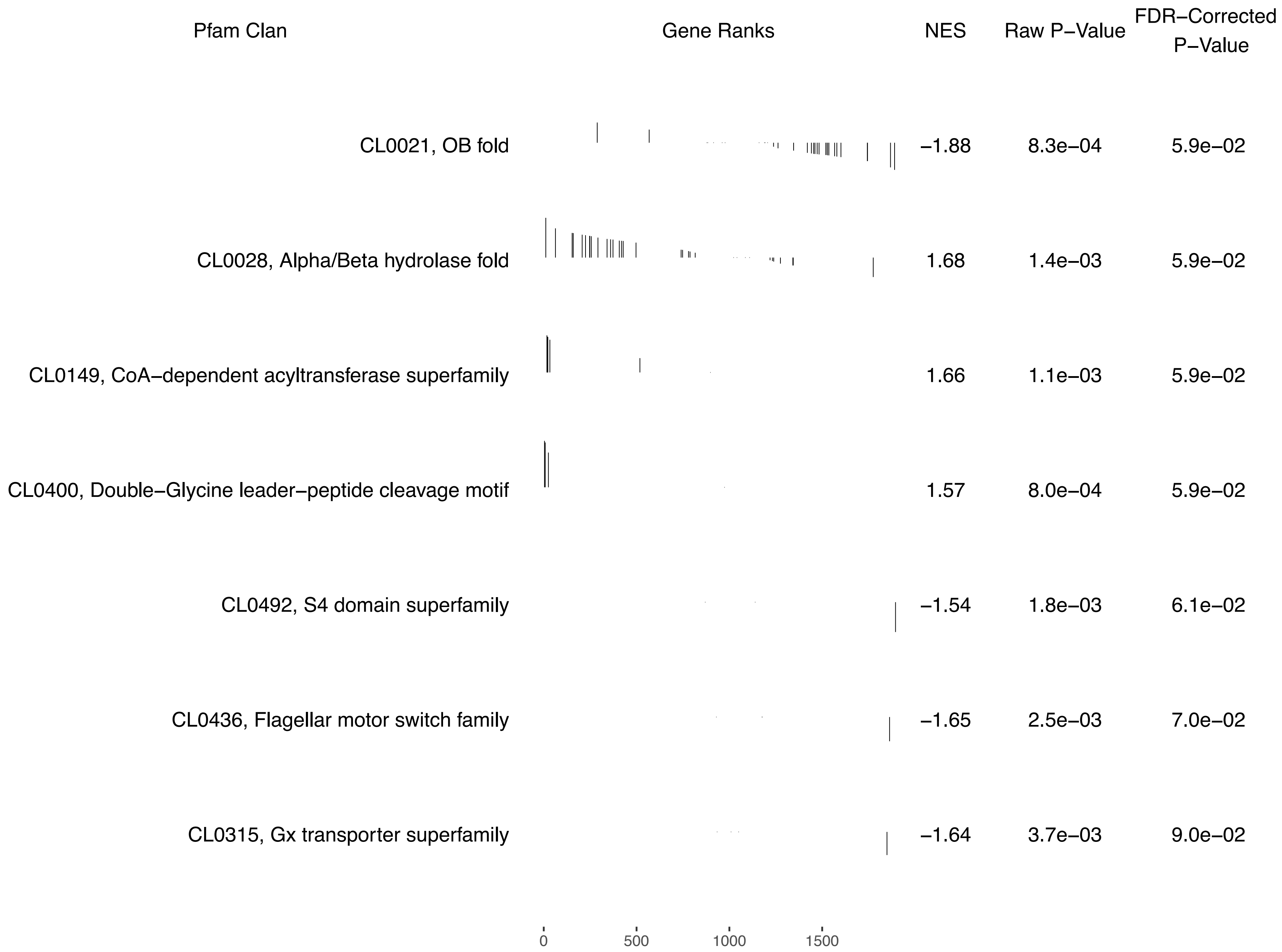


**Supplementary Figure S12.** Pfam clans (rows) enriched in BGC and non-BGC regions (false discovery rate-corrected $P < 0.10$). Normalized Enrichment Scores (NES), *P*-values, and ranks were obtained using the fgsea R package. Positive and negative NES correspond to clans enriched in BGC and non-BGC regions, respectively. Clans are ordered by false discovery rate (FDR)-corrected *P*-value (lowest-to-highest).
